# Drivers of polar bear behavior, and the possible effects of prey availability on foraging strategy

**DOI:** 10.1101/2022.07.07.498143

**Authors:** Ron R. Togunov, Andrew E. Derocher, Nicholas J. Lunn, Marie Auger-Méthé

## Abstract

Change in behavior is one of the earliest measurable responses to variation in habitat suitability, making the study of factors that promote behaviors particularly important in areas undergoing environmental change. We applied hidden Markov models to movement data of 14 polar bears, *Ursus maritimus*, from Western Hudson Bay, Canada between 2011 and 2021 during the foraging season (January–June) when bears inhabit the sea ice. The model incorporated bear movement and orientation relative to wind to classify three behaviors (stationary/drifting, area-restricted search, and olfactory search), and investigated 11 factors to identify conditions that may promote these behaviors. In contrast to other polar bear populations, we found high levels of evening activity, with active behaviors peaking around 20:00. We identified an increase in activity as the ice-covered season progressed. This apparent shift in foraging strategy from still-hunting to active search corresponds to a shift in prey availability (i.e., increase in haul-out behavior during winter to the spring pupping and molting seasons). Last, we described spatial patterns of distribution with respect to season and ice concentration that may be indicative of variation in habitat quality and segregation by bear age that may reflect competitive exclusion. Our observations were generally consistent with predictions of the marginal value theorem, and differences between our findings compared to other populations could be explained by variation in regional or temporal variation in resource abundance or distribution. Our findings and novel methodology can help identify periods, locations, and environmental conditions representing critical habitat.

## 2 Introduction

Animals exhibit a broad diversity of behaviors to meet their needs for survival, growth, and reproduction. Each behavior has consequences to the individual and has distinct relationships with the external environment (Picardi et al., 2022; Whittington et al., 2022). Changes in behavior and their associated movement patterns may represent the earliest measurable response to variation in habitat suitability and potential effects of environmental change (Wong and Candolin, 2015; Buchholz et al., 2019). For example, optimal foraging theory and the marginal value theorem predict that time spent in a resident state increases with patch quality (Charnov, 1976; Pyke et al., 1977; Nonacs, 2001). Understanding the spatial and environmental determinants of animal behavior is central to ecology and conservation. Identifying the factors that promote different behaviors may be particularly important in areas experiencing rapid environmental change or where animals occupy the limits of their realized niche. In these areas, shifts in occupied states or time budgets may precede other indicators of habitat quality (e.g., body condition, reproductive success, or survival; Wong and Candolin, 2015; Buchholz et al., 2019; Cunningham et al., 2021).

The Arctic is warming at several times the global average, resulting in reduced sea-ice extent and a prolonged ice-free period (Laidre et al., 2015b; Kwok, 2018; Onarheim et al., 2018; Stroeve and Notz, 2018; Schweiger et al., 2021). The reductions in sea ice have caused a shift toward smaller primary and secondary producers (Yun et al., 2015; Dalpadado et al., 2016), and negatively affected Arctic fish (Nahrgang et al., 2014; Christiansen, 2017) and pinnipeds (Iacozza and Ferguson, 2014; Stenson and Hammill, 2014; Hamilton et al., 2015). Polar bears (*Ursus maritimus*) rely on sea ice as a platform to access their primary prey, ringed seals (*Pusa hispida*) and bearded seals (*Erignathus barbatus*), as well as for reproduction and travel (Stirling and Archibald, 1977; Stirling et al., 2016; Florko et al., 2020b; Galicia et al., 2020). Reduction in sea ice has led to increased the energetic cost of travel. Greater habitat fragmentation has increased polar bear path tortuosity (Biddlecombe et al., 2021), more open water has increased the frequency of long-distance swimming events (Pilfold et al., 2016; Pagano et al., 2020), and increased ice drift speed has increased the cost of station-keeping (Mauritzen et al., 2003; Auger-Méthé et al., 2016; Durner et al., 2017). Further, polar bears have exhibited shifts in distribution (Lone et al., 2018), reduced access to prey (Stirling et al., 2008; Ware et al., 2017; Florko et al., 2020b), a longer fasting period (Rode et al., 2021), increased exposure to zoonotic pathogens (Pilfold et al., 2021), higher levels of cortisol (Boonstra et al., 2020), reduced body condition (Stirling and Parkinson, 2006; Rode et al., 2021), reduced access to denning habitat (Rode et al., 2021; Merkel and Aars, 2022), reduced reproduction (Stirling et al., 1999), and consequently reduced abundance in several populations (Regehr et al., 2007; Lunn et al., 2016; Regehr et al., 2016; Obbard et al., 2018; Bromaghin et al., 2021). Many of the effects of climate change on polar bears are associated with behavioral shifts including changes in foraging (Galicia et al., 2016), migration (Pilfold et al., 2016; Pagano et al., 2020), and denning strategies (Olson et al., 2017; Escajeda et al., 2018).

Polar bears exhibit a diversity of behaviors to successfully exploit the spatiotemporally dynamic sea-ice habitat (Fig. 1; Stirling, 1974; Pagano et al., 2017; Ware et al., 2020). In areas of seasonal sea ice, polar bears migrate between the terrestrial refugia and on-ice foraging grounds (Cherry et al., 2013; Bohart et al., 2021). They exhibit philopatry to their summering grounds and compensate for sea ice motion in their navigation and for station keeping (Mauritzen et al., 2003; Auger-Méthé et al., 2016; Durner et al., 2017; Klappstein et al., 2020). Polar bears rely both on visual and olfactory search to hunt sparsely distributed prey (Stirling, 1974; Smith, 1980), which may be influenced by presence of daylight (Togunov et al., 2017, 2018). However, the expansive and remote nature of their habitat impedes behavioral research. Direct observational research (e.g., Stirling, 1974; Jagielski et al., 2022) is limited in spatial or temporal extent. Insight into polar bear ecology across larger spatiotemporal scales often relies on remote tracking data, however these studies are typically not behaviorally-explicit (e.g., Kolz et al., 1980; McCall et al., 2016; Laidre et al., 2018; Durner et al., 2019). Although there have been a few large-scale multi-behavioral studies using polar bear telemetry (e.g., Auger-Méthé et al., 2016; Pagano et al., 2020; Ware et al., 2020), these were either limited to two simple behavioral states (e.g., active or inactive) or did not investigate associations between behaviors and the environment.

**Figure 1:**
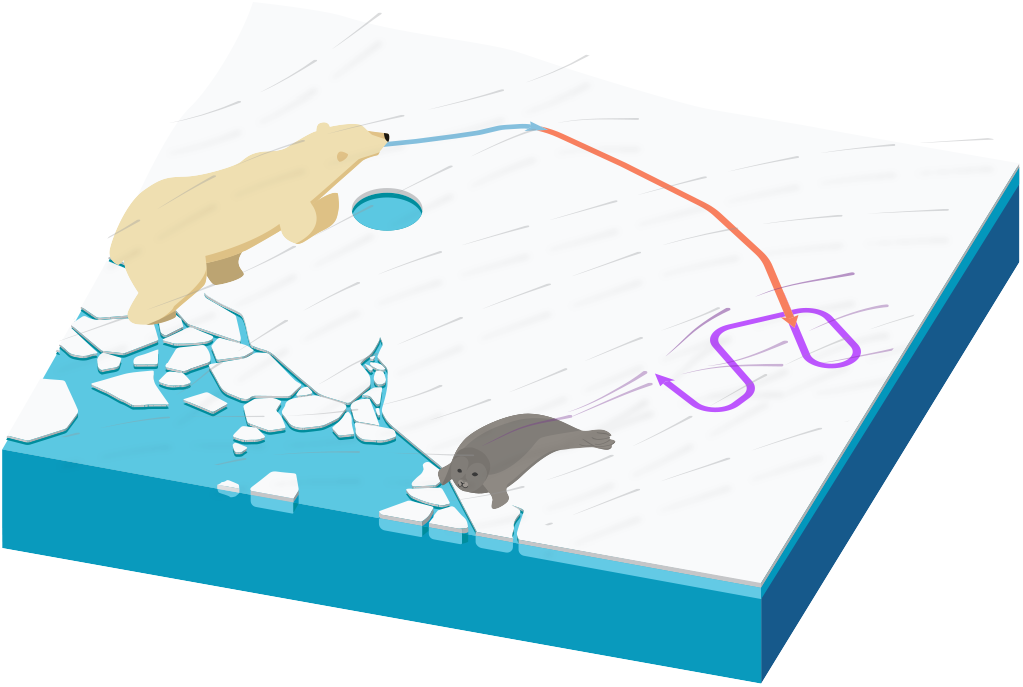
Illustration of polar bear behavior and movement. Polar bear track drifting ≈ 15° relative to wind (gray) when stationary on sea ice (e.g., when still-hunting by breathing hole; blue), moving ≈ 90° relative to wind during olfactory search (red) to maximize probability of encountering odor plumes (purple), and random movement relative to wind during area-restricted search (purple track). Reproduced with permission from (Togunov et al., 2021).

Recent advances in data acquisition (Nathan et al., 2022) and analytical methods (e.g., McClintock and Michelot, 2018; Togunov et al., 2021) have enabled the identification of more intricate behaviors and research at an increasingly large scale and resolution. Pagano et al. (2017) described the use of accelerometers to identify up to ten fine-scale behaviors, and Pagano et al. (2020) used a combination of accelerometer data and conductivity sensors to identify resting, walking, and swimming. Unfortunately, most existing telemetry datasets do not lend themselves to many of the newer analytical methods because they lack necessary auxiliary data (i.e., they only estimate tag location). However, Togunov et al. (2021) described the use of location data in combination with wind to identify up to three behavioral states, a method readily applicable to most existing polar bear movement data.

Using remote tracking data for behavioral research is further complicated by sea ice motion. The motion of sea ice is imparted on the polar bear track, changes the apparent speed and orientation, and complicates behavior characterization and classification (Gaspar et al., 2006; Auger-Méthé et al., 2016; Togunov et al., 2021). The conventional method used to remove sea ice motion is to subtract satellite-based estimates of ice velocity from the movement track (e.g., Auger-Méthé et al., 2016; Blanchet et al., 2020; Klappstein et al., 2020), however, this approach assumes there is negligible bias in the sea ice motion data, which is typically not the case (Dohan and Maximenko, 2010; Yonehara et al., 2016; Togunov et al., 2020). Togunov et al. (2021) described the use of biased correlated random walk (BCRWs) models to identify periods where the bear is stationary and passively drifting on the sea ice (Fig. 1). However, non-stationary behaviors may still be misclassified. Therefore, there is a need for movement models that can correct for sea ice motion that are flexible to error in environmental data.

Given climate change induced alterations of sea ice, we were interested in understanding how polar bears change their behavior in space and time and how these behaviors were affected by environmental variability. Our objectives were to: (1) develop a sea ice motion correction model that account for error in satellite-based estimates, (2) examine behavior time budgets during the winter foraging period, (3) identify factors associated with different behaviors, and (4) examine broad-scale and behavior-specific habitat use.

## 3 Methods

### 3.1 Study area and telemetry data

Hudson Bay, Canada is a large inland sea with an area of 830,000 km^2^ (Fig. 2; Prinsenberg, 1984). The Bay is characterized by seasonally-present sea ice, which begins to form during freeze-up in late November. The ice covers the majority of the Bay from January to May and is comprised primarily of drifting pack ice and a 10-15 km wide fringe of land-fast ice (Lamont, 1948; Canadian Coast Guard, 2012). The sea ice concentration begins to decline in May, typically reaching 50% cover in early July, and melting completely by early August (Danielson, 1971; Saucier et al., 2004). The focal time period of this study was January to July, representing the primary feeding and mating period and the subsequent decline in sea ice (i.e., break-up).

**Figure 2:**
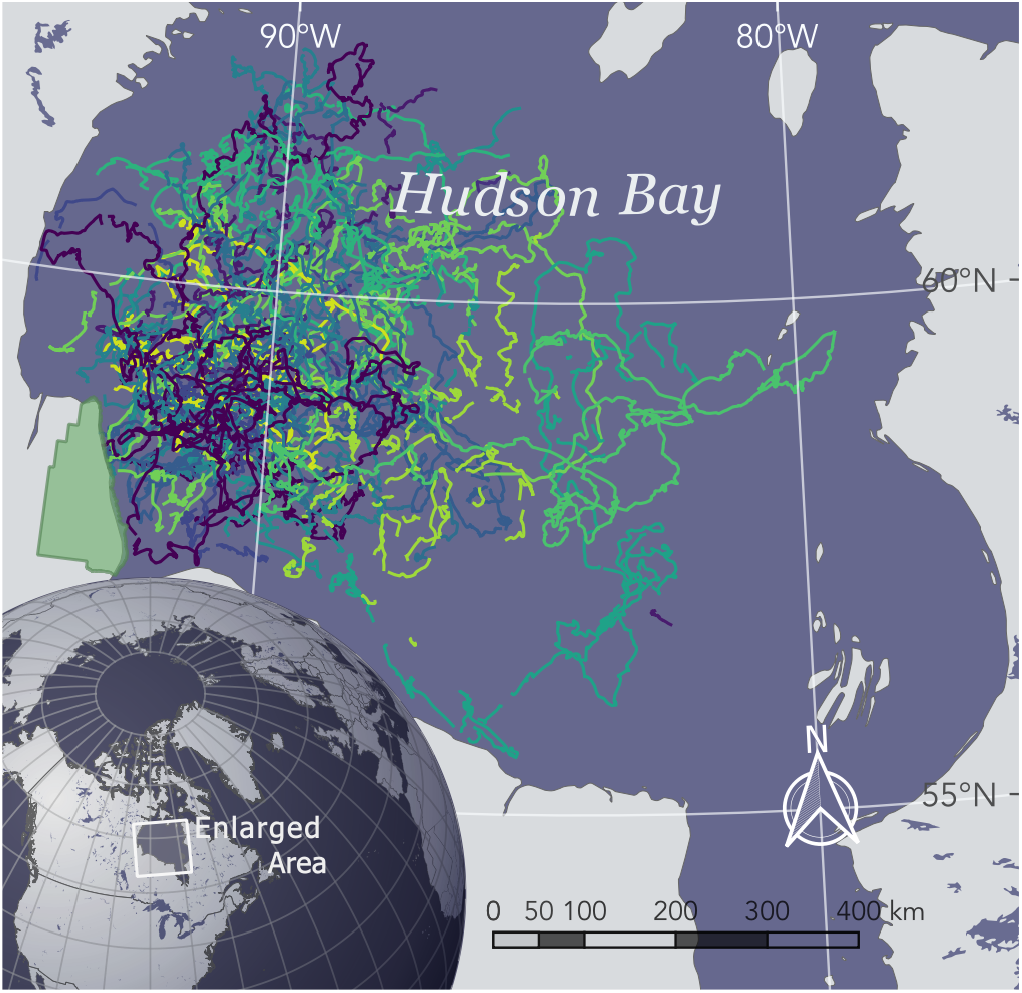
Hudson Bay study area (enlarged) and polar bear tracks (Colored lines). Gaps in telemetry data > 6 h and bouts *<* 24 h were excluded from analysis and are not shown. The green region represents Wapusk National Park.

As part of a long-term study of the population ecology of polar bears in Western Hudson Bay (Ramsay and Stirling, 1988; Stirling et al., 1999; Regehr et al., 2007; Lunn et al., 2016), 107 adult females with cubs were tranquilized from helicopters (Stirling et al., 1989) during the summers of 2010 – 2019 and equipped with Argos® or Iridium® satellite-linked global positioning system (GPS) collars (Telonics, Mesa, AZ). Lone females were not collared as they may have been pregnant and would remain in maternity dens up to seven months after collaring. Males were not collared because their neck circumference is greater than that of their head and would not retain collars. Tagging was performed primarily in the Western Hudson Bay polar bears’ summering and denning grounds in Wapusk National Park, Manitoba (Fig. 2; Stirling, 1977; Richardson et al., 2005; Florko et al., 2020a). A vestigial premolar was extracted from each collared bear to determine age by counting annuli in the cementum (Calvert and Ramsay, 1998). The animal handling protocols performed were approved by the University of Alberta Animal Care and Use Committee for Biosciences and by the Environment Canada Prairie and Northern Region Animal Care Committee (permit numbers: AUP00000033 and AUP00003667).

The collars were programmed to last two years after which they would release on a predefined date. Of the 107 collars deployed, 79 obtained locations every 4 h, 25 collars obtained locations every 2 h, and three collars obtained locations every 30 min. A preliminary analysis indicated that the 4-h collars could reliably only identify two behaviors (data not shown). To identify at least three behaviors, we used only the collars at 2-h or 30-min frequency, and we sampled the 30-min data to a 2-h frequency. The bear locations were projected into a Universal Transverse Mercator coordinate reference system (NAD83 Teranet Ontario Lambert, EPSG: 5321).

We used hidden Markov models (HMM) to investigate the relationship of different behavioral states in relation to environmental covariates. The classic HMM assumes the observation data is in discrete time and that there is no missing data in the predictor variables (Zucchini et al., 2016; McClintock and Michelot, 2018). To avoid interpolating large gaps, we segmented the location data into separate bouts whenever missing locations spanned more than 6 h (i.e., we interpolated a maximum of three missing locations). To remove data-sparse bouts, we removed segments spanning less than 24 h or those with fewer than eight locations (as in Togunov et al., 2021). Any missing locations in the remaining bouts were interpolated using the R Package `crawl` (Johnson et al., 2008; Johnson and London, 2018). The remaining telemetry data used in the subsequent analysis are presented in Fig. 4.

**Figure 3:**
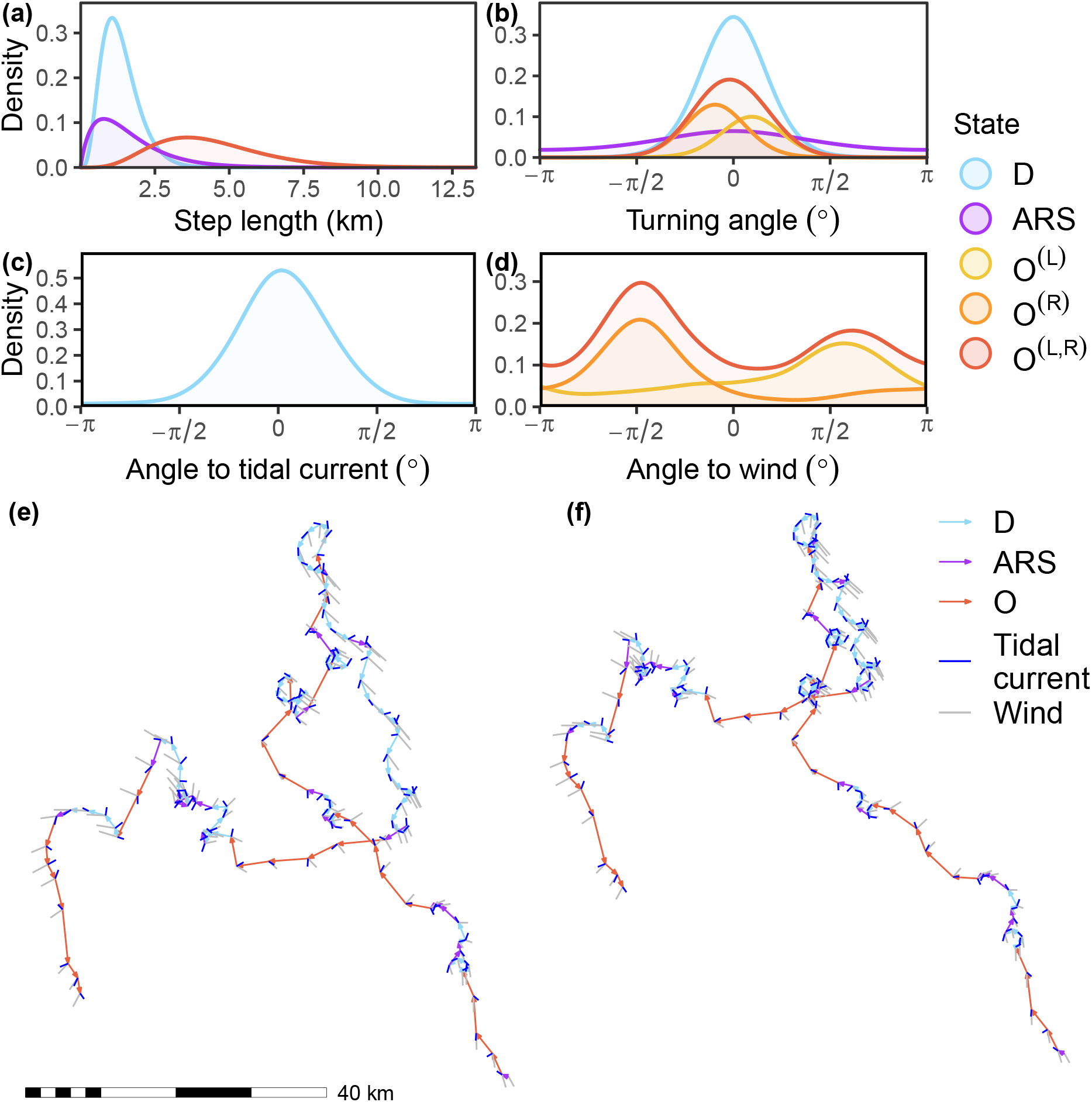
Predicted state characteristics. (a) Step length distribution for each state; (b) turning angle distribution for each state; polar bear orientation relative to (c) tidal currents and (d) wind; and example from one bear bout contrasting the (e) original track and (f) wind-forcing-corrected track. *D* and *ARS* represent drift and area-restricted search, respectively, and *O* represent olfactory search (left, *O*^(*L*)^, or right, *O*^(*R*)^, relative to wind). Tracks in (e) and (f) are colored by the decoded states and show the estimated wind (gray) and tidal current (blue) velocities. All data (except in panel (e)) were based on the wind-forcing-corrected tracks and the top model fitted to seven years of polar bear telemetry data from Western Hudson Bay, Canada.

**Figure 4:**
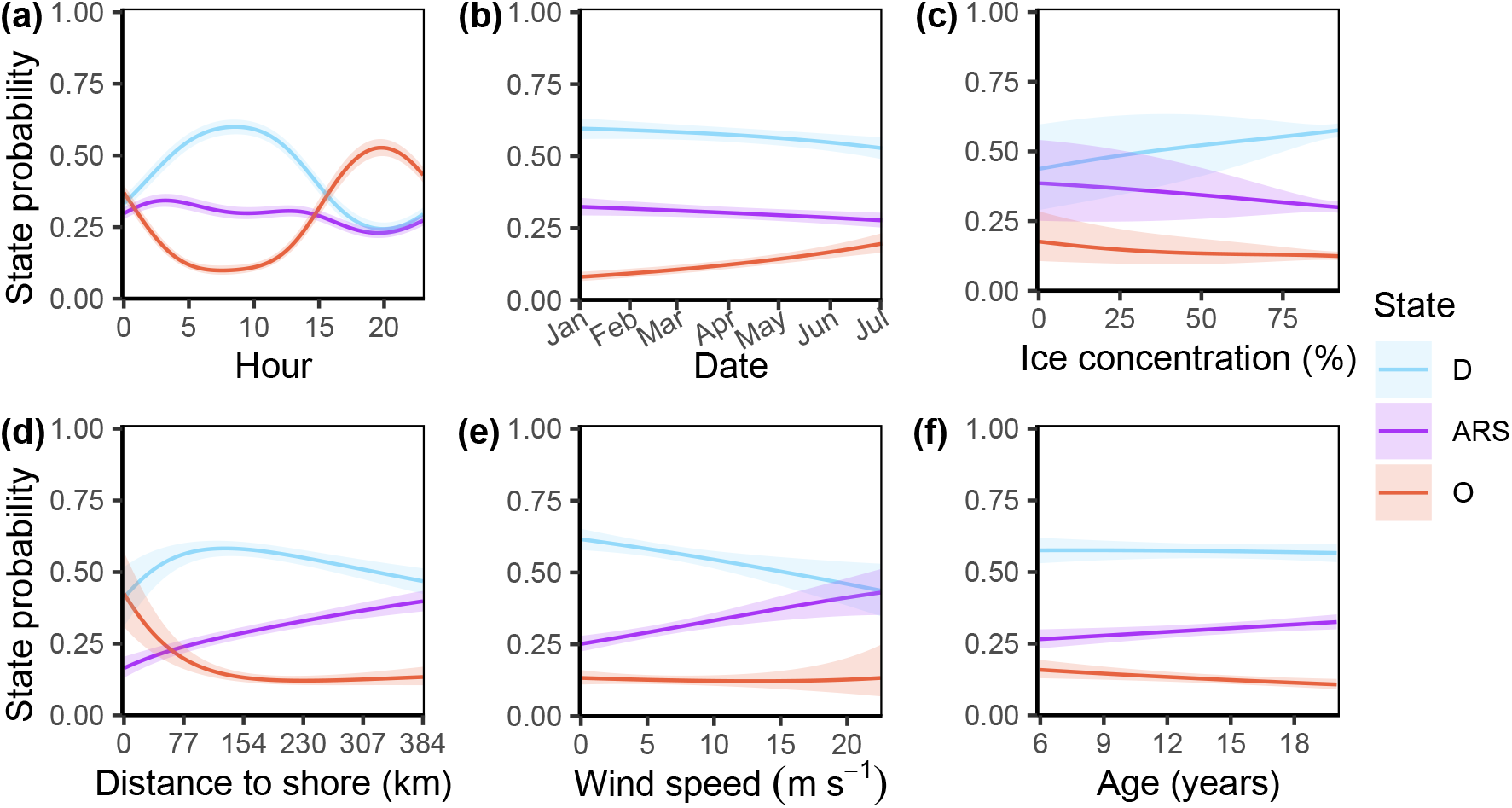
Stationary state probabilities as functions of (a) hour, (b) day, (c) ice concentration, (d) distance to shore, (e) wind speed, and (f) bear age. Shaded areas represent the 95% confidence interval. *D, ARS*, and *O* represent drift, area-restricted search, and olfactory search, respectively.

Next, we annotated this telemetry data with the following covariates: bear age and cub status from when the bears were tagged; ordinal date and hour of the GPS fix; sun altitude calculated using the R package `oce` (Kelley et al., 2022); wind velocity, snow depth, total precipitation, surface solar radiation from the ERA5 meteorological reanalysis project (Hersbach et al., 2020); sea ice concentration (Comiso, 2017); bathymetry (IOC et al., 2003); tidal currents (St-Laurent et al., 2008); and sea ice motion vectors by the National Snow and Ice Data Center (NSIDC; Tschudi et al., 2019, 2020). Spatial covariates provided as a gridded raster were interpolated in space and time as in Togunov et al. (2017, 2018).

### 3.2 Sea ice motion correction

To correct for sea ice motion in the telemetry data, without assuming the environmental data is free of bias, we first fitted a BCRW model to tracks of dropped collars (Togunov et al., 2020). This BCRW described the motion of the dropped collars as a function of either ERA5 wind velocities (Hersbach et al., 2020) or NSIDC sea ice motion vectors (Tschudi et al., 2019, 2020). The data source (i.e., ERA5 wind or NSIDC sea ice motion) that predicted sea ice motion was used for the correction. In this instance, the hourly wind data outperformed the daily NSIDC data (details in Supplementary materials Appendix B). Next, we used the BCRW fitted to the dropped collars to predict and subtract the estimated component of sea ice motion driven by wind from the bear telemetry data (details in Supplementary materials Appendix A). After correcting for wind-advection, the 2-h bear collars still retained motion from tidal currents, which complete 360° counter-clockwise rotations approximately every 12 h (St-Laurent et al., 2008). This tidal motion could not be corrected using the dropped collars due to their lower 4-h resolution (Togunov et al., 2020). Therefore, the residual tidal current was integrated into the motion of the drift state in the behavioral model (details in Section 3.3). Although it is challenging to remove sea ice motion from the movement track, we expected reducing its effect would increase the accuracy of the HMM – particularly for characterizing states with movement speed similar to sea-ice speed.

### 3.3 Behavior analysis

Some behaviors exhibit orientation bias relative to the external stimuli, which is defined as *taxis* (Codling et al., 2008). The degree of bias can occur along a spectrum from being primarily governed by bias relative to external stimuli (biased random walk; BRW; all notation used in this paper is described in Table 1), to a trade-off between directional persistence and orientation bias (biased correlated random walk; BCRW), to being primarily governed by directional persistence (CRW; Benhamou, 2006; Codling and Bode, 2016; Codling et al., 2008). Bias toward an angle relative to a stimulus is defined as *menotaxis*.

**Table 1:**
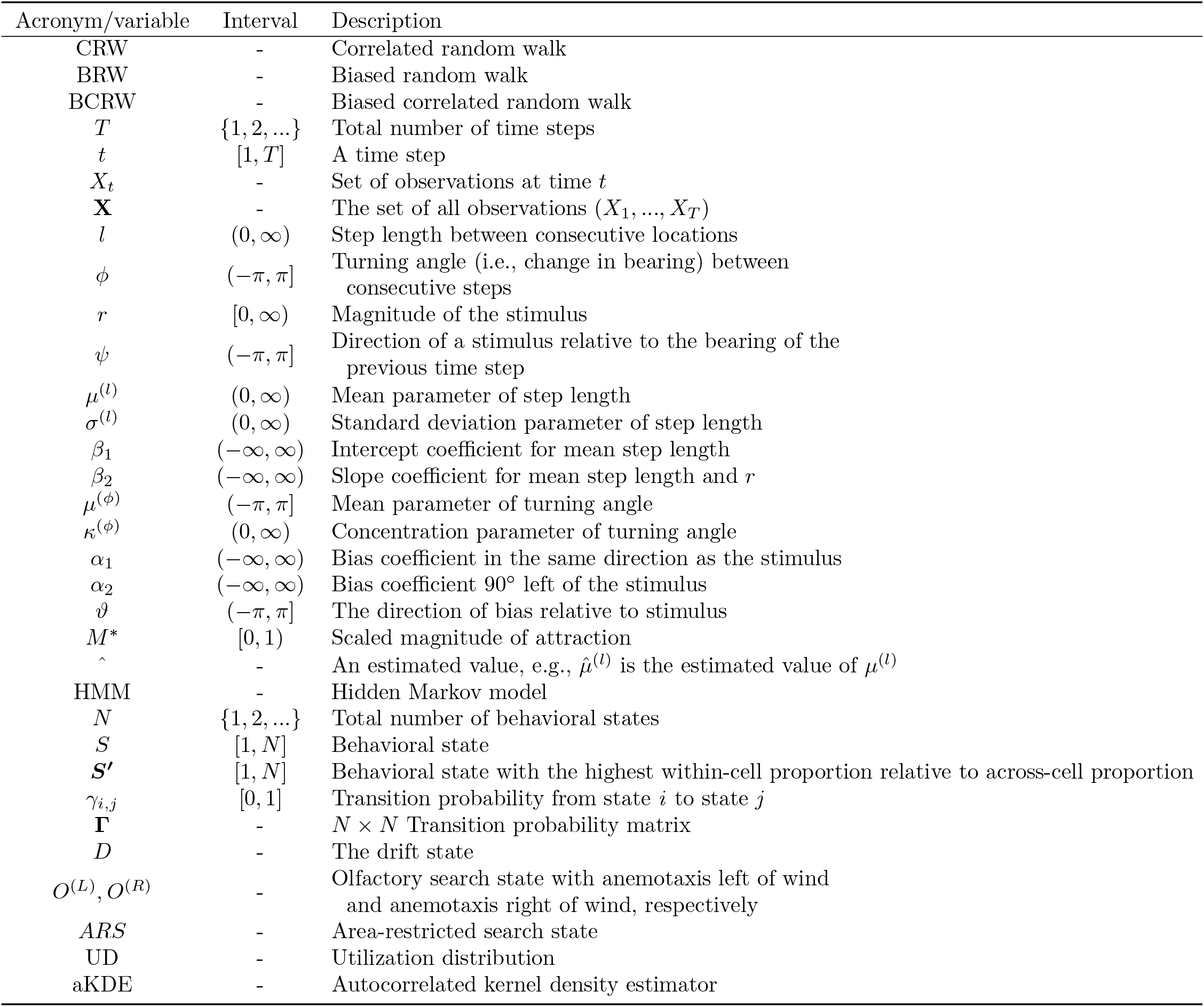
Description of acronyms and notation used in this paper and their interval, if applicable.

We used an adapted version of the HMM with menotactic behaviors described in Togunov et al. (2021) to investigate three movement behaviors: drifting, area-restricted search (*ARS*), and olfactory search. We described these behaviors using a four-state HMM in which the drifting and *ARS* behaviors were represented by their own respective states (*D* and *ARS*) and olfactory search was divided into two discrete states corresponding to bias left and bias right relative to wind (*O*^(*L*)^ and *O*^(*R*)^, respectively; Fig. 1; Togunov et al., 2021). Under this HMM framework, the states were assumed to be a discrete-time latent Markovian process, where the probability of a state *S*_*t*_ at time *t* ∈ 1, …, *T* depends only on the state at the previous time *S*_*t*−1_, and the observed data **X**_*t*_ depends only on the hidden state *S*_*t*_ (Zucchini et al., 2016; McClintock et al., 2020). The state transition probabilities *γ*_*i,j*_ = Pr(*S*_*t*+1_ = *j*|*S*_*t*_ = *i*) for *i, j* ∈ {1, …, *N*} (where *N* is the number of states) are summarized by the *N* × *N* transition probability matrix, **Γ**. We extracted two variables from the tracking data: step length *l*_*t*_ ∈ (0, ∞) (the distance between consecutive locations) and turning angle *ϕ*_*t*_ ∈ (−*π, π*] (change in bearing between consecutive steps; Langrock et al., 2012; McClintock and Michelot, 2018; Togunov et al., 2021). Following Togunov et al. (2021), we assumed step lengths followed a gamma distribution:

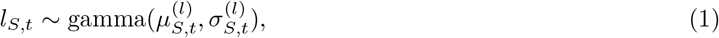

where 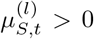 and 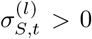 are the state-specific mean and standard deviation, respectively, of the step length at time *t* (McClintock and Michelot, 2018; Togunov et al., 2021). The motion of the drift state was defined as a function of predicted tidal currents as it could not be corrected for using the 4-h dropped collars. Specifically, we defined the mean step length following:

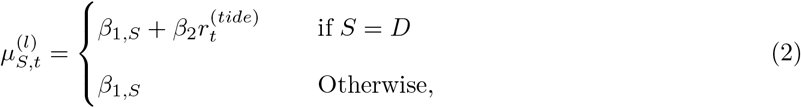

where *β*_1,*S*_ ∈ (−∞, ∞) is the state-specific intercept coefficient for step length mean and *β*_2_ ∈ (−∞, ∞) is a slope coefficient representing how the mean step length of *D* is affected by tidal speed 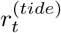. Further, we used pseudo-design matrices in conjunction with working boundaries to ensure that the step length of olfactory search was faster than ARS and that ARS was faster than drift (i.e., 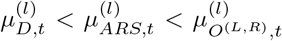; details in Supplementary materials Appendix C; McClintock and Michelot, 2018).

We assumed the turning angle followed a von Mises distribution:

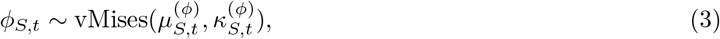

where 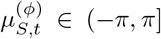 is the state-specific mean turning angle parameter at time *t* and 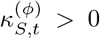 is the state-specific concentration parameter around 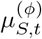 (McClintock and Michelot, 2018; Togunov et al., 2021). We assumed the mean turning angle for *D* was a circular-circular regression function of tidal currents and that the degree of bias toward the direction of tides increased with tidal speed (details in Supplementary materials Appendix C). The *ARS* state turning mean angle was fixed at 0 (i.e., 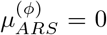). The olfactory search states were modeled as menotactic BCRWs with biases toward a unknown angles relative to wind. Following Togunov et al. (2021), this menotactic BCRWs modeled the mean turning angle as a trade-off between bias parallel to wind and bias perpendicular to wind. The mean turning angle of each behavior was modeled as follows:

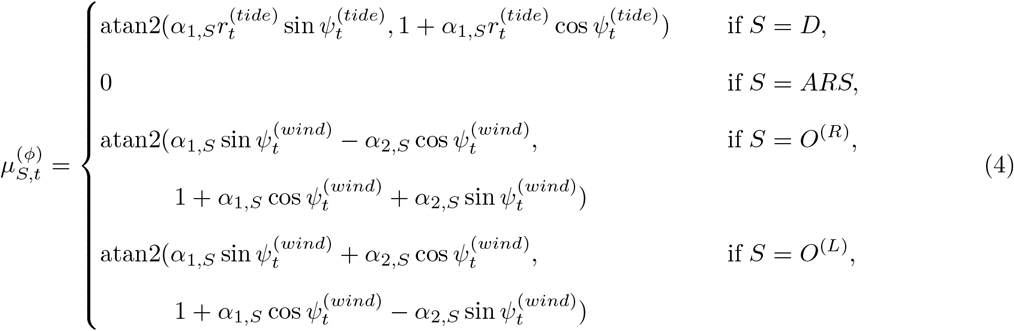

where 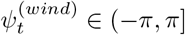 and 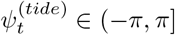 represent the directions of wind and tides, respectively, at time *t* relative to the track bearing at time *t* − 1, *α*_1,*S*_ ∈ R represents the state-specific bias coefficient parallel to *ψ*_*t*_, and *α*_2,*S*_ ∈ (−∞, ∞) ∈ R represents the state-specific bias coefficient toward 90° anti-clockwise of *ψ*_*t*_ (Togunov et al., 2021). As *O*^(*L*)^ and *O*^(*R*)^ represented the same underlying behavior, they shared the parameters for state transition probabilities *γ*_*i,j*_, step length 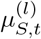 and 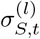, turning angle concentration 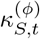, and bias parallel to wind *α*_1,*S*_, and we fixed *α*_2,*O*(*R*)_ = *α*_2,*O*(*L*)_ (Togunov et al., 2021). Following Togunov et al. (2021), the angle of attraction relative to wind was represented by *ϑ* = atan2(*α*_2_, *α*_1_). In addition, we could represent where a behavior lies along the spectrum of CRW and BRW using the scaled magnitude of attraction 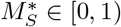:

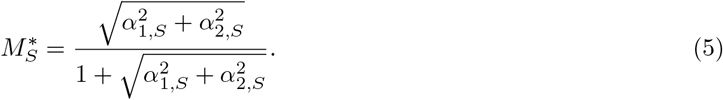

The CRW and BRW are limiting cases of Eq. 5, where *M*^*^ → 0 and *M*^*^ → 1, respectively, while a BCRW would have an intermediate value of *M*^*^. That is, a value of *M*^*^ close to 0 represents behavior with high directional persistence, and a value close to 1 represents behavior with orientation highly correlated to an external stimuli.

To investigate how environmental conditions influence the probability of different behaviors, we modeled the state transition probabilities as functions of the annotated covariates (Langrock et al., 2012; McClintock and Michelot, 2018). To allow for non-linearity, ordinal date, sun altitude, wind velocity, surface solar radiation, ice concentration were fitted in linear and quadratic form, and hour of the day was fitted as a cosinor model (McClintock and Michelot, 2018). To determine which form (i.e., linear or quadratic, cosinor) to use, we fitted HMMs with each form of the covariate on their own (i.e., with no other covariates), and selected the form with the lowest Akaike information criterion (AIC; Akaike, 1974). Next, we used forward and backward AIC model selection to determine which combination of covariates best explained state transitions (Patterson et al., 2017; Byrnes et al., 2021).

Over time, state probabilities of a Markov chain converge to the ‘stationary distribution’, which represent the marginal probability of a state assuming the covariates remains constant (Patterson et al., 2009; Langrock et al., 2012; Zucchini et al., 2016). We estimated the 95% confidence interval on the stationary distribution as an indicator of significant within-state change in relation to covariates in state probability or between-state probability McClintock and Michelot (2018). Because we were not examining variation between bears, we did not include random effect on individual ID, and all estimated model coefficients were shared among individuals (McClintock and Michelot, 2018; McClintock, 2021). The HMMs were fitted using the R package `momentuHMM` (McClintock and Michelot, 2018).

To investigate the spatial distribution of states, we first determined the most likely state for each step using the Viterbi algorithm (Zucchini et al., 2016). Second, we rasterized the state-decoded steps by assigning them to cells of a regular 50 km grid. Third, to reduce temporal autocorrelation, we rarefied the steps into unique “bear days”, such that for each cell, we retained only one data point for unique bear ID, date, and state. A simple measure of the most frequent state in a cell would bias states that were more common overall and fail to identify where uncommon states were disproportionately frequent. Therefore, for each cell, we identified which state was most frequent relative to the frequency of each state across all cells. Specifically, for each cell *i*, we defined the state ***S′*** as the state with the highest within-cell proportion relative to proportion across all cells following:

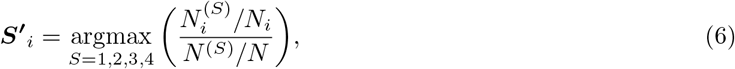

where 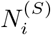 is the number of bear days in cell *i* for state *S, N*_*i*_ is the total number of bear days in cell *i* across all states, *N* ^(*S*)^ is the number of bear days across all cells for state *S*, and *N* is the total number of bear days across all cells and states. ***S′*** was calculated for the entire data set, as well as separately for early winter (January–March) and late winter (April–June) to compare seasons, separately for years with low ice concentration and high ice concentration to compare years with different conditions, and separately for younger (6–14 years) to older (15–20 years) adult bears to compare age classes. We obtained weekly sea-ice coverage in Hudson Bay from January to June from Ice Graph version 2.5 (Canadian Ice Service, 2021). We calculated the mean total ice concentration for each year and classified each into either years with below or above average mean total ice concentration. Spatial state segregation was described qualitatively. In addition to spatial state segregation, we compared the spatial extent of telemetry data by season and between years with high and low concentration. Utilization distribution (UD) was calculated using the 80% autocorrelated kernel density estimator (aKDE) using the R package `ctmm` (Calabrese et al., 2016); this analysis was only descriptive and did not assess statistical significance. All analyses were conducted in `R` version 4.1.1 (R Core Team, 2020).

## 4 Results

We obtained 31550 locations during the focal months of January to June from 14 unique bears. Some collars provided data across multiple years, yielding 24 unique “bear-years”. The movement bouts contained 2484 (7%) missing locations that were interpolated using the R package `crawl`. After filtering out short and data-sparse bouts from the interpolated segments, we retained 33160 unique steps. Each year had between 2084 (2019) and 10092 (2020) locations, and we obtained between 4199 (February) and 6545 (April) locations across months.

Both forward and backward model selection converged to the same top model. In this model, drift was characterized as a slow BCRW with a moderate turning bias in the same direction as tidal currents. At the median estimated tidal speed of 0.29 km h^−1^, the mean step length of drift was 0.63 ± 0.31 km h^−1^ 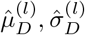; Table 2, Fig. 3a). The turning angle concentration (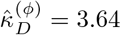; Table 2, Fig. 3b) and scaled magnitude of attraction (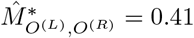; Table 2) were moderate, best characterizing drift as a BCRW. The direction of tidal currents explained much of the variation in direction of drift with the majority of drift direction falling within 5.0° ± 43.8 SD of the tidal currents (Fig. 3c and f).

**Table 2:** Parameter estimates for four-state HMM.

| State | $\hat{\mu}^{(l)\dagger}$ | $\hat{\sigma}^{(l)\ddagger}$ | $\hat{\vartheta}^{\S}$ | $\hat{\kappa}^{(\phi)\P}$ | $\hat{M}^{*\parallel}$ |
| --- | --- | --- | --- | --- | --- |
| Drift, $D$ | 0.63 | 0.31 | NA | 3.64 | 0.41 |
| Area-restricted search, $ARS$ | 0.76 | 0.59 | NA | 0.62 | NA |
| Olfactory search, $O^{(L,R)}$ | 2.04 | 0.93 | 92 | 4.28 | 0.29 |

† mean step length ( $\text{km h}^{-1}$ ), ‡ step length standard deviation ( $\text{km h}^{-1}$ ), $\S$ angle of attraction relative to wind ( $^\circ$ ), $\P$ turning angle concentration, $\parallel$ scaled magnitude of attraction.

ARS was characterized as a slow CRW with no bias relative to wind. The estimated mean step length was 0.76 ± 0.59 km h^−1^ (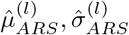; Table 2, Fig. 3a). Mean turning angle was fixed to zero and the turning angle concentration was the lowest among the states (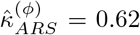; Table 2, Fig. 3b), best characterizing ARS as a CRW with low persistence.

Olfactory search was characterized as a fast BCRW with a bias relative to wind. Olfactory search had the highest estimated mean step length (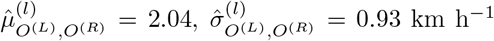; Fig. 3a), with an overall bias toward ±92° relative to wind (downwind bias of 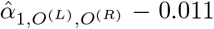, and crosswind bias of 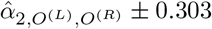; Table 2, Fig. 3d). The turning angle concentration was the highest among the states (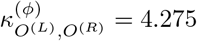; Table 2, Fig. 3b). The scaled magnitude of attraction relative to wind was moderately 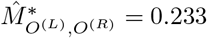 Table 2), best characterizing olfactory search as a BCRW.

Based on the Viterbi-decoded states, polar bears spent approximately 47% of their time in the drift state, 24% in ARS, and 29% in olfactory search. Six covariates on the state transition probability were identified in the top model: hour, day, ice concentration, distance to shore, wind speed, and bear age. The highest variability in state probability was with respect to hour of the day. The drift state was most frequent between 00:00 h and 15:30 h (local time, UTC -5 H) with a peak around 08:30 h. Olfactory search was most frequent between 15:30 and 00:00 with a peak around 20:00 h. However, this evening peak activity appeared to be primarily driven by movement in January–May, while bears did not appear to exhibit significant diurnal variation in June (Supplementary materials Fig. D1). ARS exhibited little variation compared to drift and olfactory search, though it appeared to decrease from a peak at 03:00 h to a trough at 20:00 h (Fig. 4a). With respect to ordinal date, drift and ARS decreased as the season progressed, while olfactory search increased (Fig. 4b). With respect to ice concentration, the confidence intervals decreased with increasing ice concentration, likely due to the larger amount of data at higher concentrations. ARS and olfactory search marginally decreased with increasing concentration, while drift increased with ice concentration. At ice concentrations *<* 50%, drift and ARS had similar probabilities (Fig. 4c). The probability of ARS gradually increased as distance to shore increased. The probability of being in a drift state increased up to ∼ 130 km from shore, then gradually declined. The probability of olfactory search declined rapidly until a distance of ∼ 150 km from shore, then remained relatively consistent. Near shore, the probabilities of drift and olfactory search were similar, and at a maximum distance of ∼ 390 km, the probabilities of drift and ARS were similar (Fig. 4d). As wind speed increased, the probability of drift decreased, ARS increased, while olfactory search remained relatively consistent. At the highest observed wind speeds ∼ 20 m s^−1^, the probabilities of drift and ARS were similar (Fig. 4e). As age increased, the probability of ARS increased, olfactory search decreased, while drift remained relatively consistent (Fig. 4f).

There was a non-uniform distribution of location data across the Bay, with the highest concentration of locations occurring around −90.5° longitude and 58.5° latitude (Fig. 5a). There appeared to be marginal segregation of states, with drift appearing to be more common west of −89° longitude, ARS was more common east of −89° longitude, and olfactory search was more common around the periphery of the overall extent (Fig. 5b). The spatial segregation between drift and ARS was apparent in late season, when mean annual ice concentration was high, and among older bears (Fig. 6). The UD was ≈ 31% greater during early winter (January–March; 80% UD = 316,550 km^2^) compared to late winter (April–June; 80% UD = 242,100 km^2^; Fig. 6a and b). Compared to years with above average ice concentration ice concentration (2018, 2019, and 2020; *μ*±SE= 91.95 ± 0.31), the 80% UD in years with below average ice concentration (2011, 2016, 2017, and 2021; *μ*±SE= 89.03 ± 0.28) was ≈ 15% greater (low: 277,753 km^2^, high: 267,418 km^2^; Fig. 6c and d). The 80% UD of younger individuals was ≈ 5% smaller compared to older individuals (younger: 312,822 km^2^, older: 330,313 km^2^; Fig. 6e and f).

**Figure 5:**
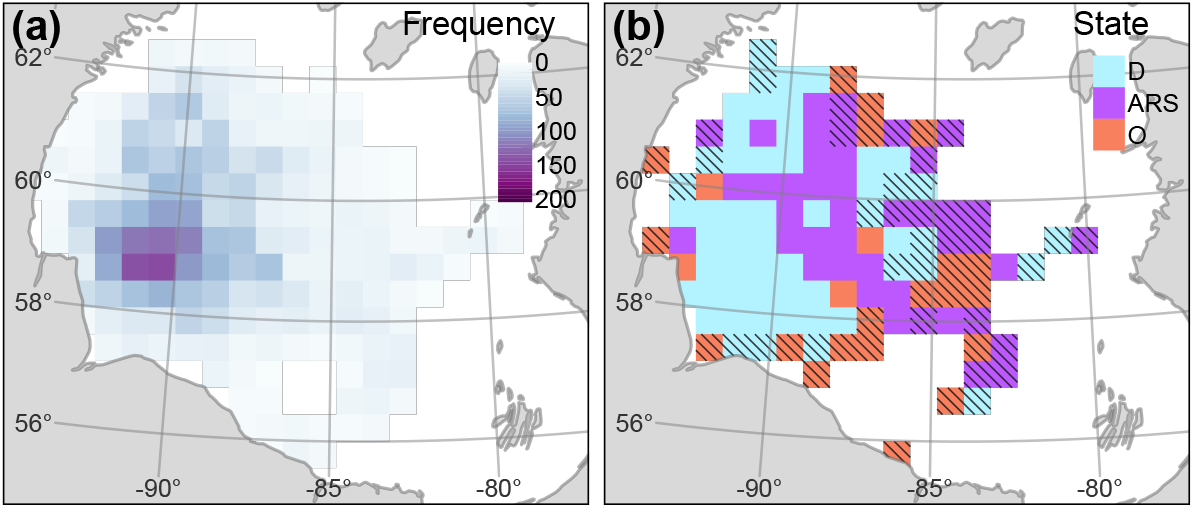
Distribution of predicted states. (a) total number of bear days and (b) state ***S′*** with the highest within-cell proportion relative across-cell proportion. Cells with *<* 7 bear days were not plotted and cells with *<* 21 bear days were hashed.

**Figure 6:**
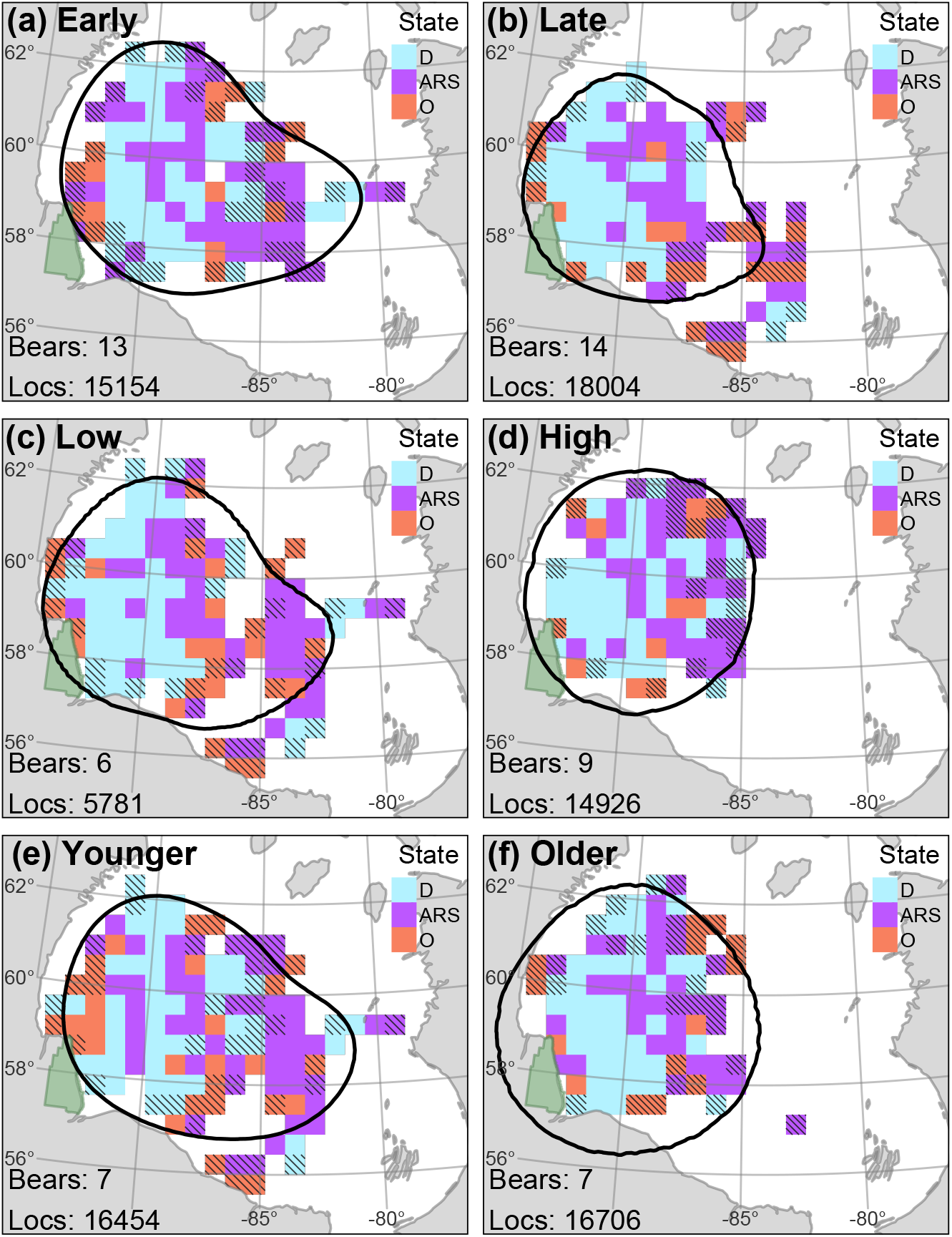
Maps of state ***S′*** with the highest within-cell proportion relative to across-cell proportion. Data were subset into (a) early winter (January–March) or (b) late winter (April–June); (c) years with low ice concentration (2011, 2016, 2017, and 2021) or (d) years with high ice concentration (2018, 2019, and 2020); and (e) youngest seven bears (6–14 years) or (f) oldest seven bears (15–20 years). Maps were based on Viterbi-decoded states and rarefied to bear days. Cells with *<* 7 bear days were not plotted and cells with *<* 21 bear days are hashed. The number of unique bears and number of locations are presented on the bottom left. The green region represents Wapusk National Park. Contour line represents the 80% utilization distribution.

## 5 Discussion

Polar bears exhibit a high degree of behavioral plasticity and diversity (Stirling, 1974; Thiemann et al., 2008; Pagano et al., 2017; Ware et al., 2020), however the remote and dynamic nature of their habitat has made it difficult to study their behavior, particularly during the critical winter foraging period. We used remote tracking data to investigate the spatiotemporal distribution of three movement states representative of three important behaviors (stationary/drifting, area-restricted search, and olfactory search) and to examine what factors may promote them. We identified six factors that appear to affect state probability and the spatial variation in state distribution and segregation. Most notably, we observed diurnal and intra-annual variation that may be indicative of a circadian rhythm and seasonal shifts in foraging strategy corresponding to known changes in prey availability. In addition, we observed variation in the spatial extent of movement that may be related to variation in habitat quality or intraspecific competitive exclusion.

One of the key challenges when applying HMMs to identify behavior from movement data is the biological interpretation of states (Pohle et al., 2017; McClintock et al., 2020). We classified movement into three states, each of which may represent more than one behavior. For example, the drift state may represent any stationary behavior, including sheltering during adverse weather, resting, still-hunting, or prey handling. To facilitate the interpretation of the results, we used prior knowledge on the behavior and phenology of polar bears and their prey. In addition, we interpreted states through the lens of classic optimal foraging theory and the marginal value theorem, which provide predictions on relationships between residency times and energy expenditure in relation to resource availability and habitat quality (Charnov, 1976; Pyke et al., 1977; Florko et al., In review).

### 5.1 State probability

Nearly half of polar bears’ overall time budget was occupied by the sedentary drift behavior, which was also the most frequent state. These results align with previous work revealing that polar bears spend the majority of their time in stationary behaviors (Stirling, 1974; Pagano et al., 2017). For the first 15 h of the day, drift was the most prevalent state with a peak around 08:30 h. Similarly, Ware et al. (2020) detected minimal activity before 08:00 h during January–March for the Southern Beaufort Sea polar bear population. Moreover, Stirling (1974) identified the first 8 h of the day as the most advantageous period for still-hunting seals, the favored hunting strategy of polar bears during the non-pupping season. Seal haul-out varies diurnally and seasonally. During early winter, seals haul-out primarily at night, while during spring, haul-out behavior peaks around 12:30 h and the highest number of seals in the water occurring at night (Stirling, 1974; Frost et al., 2004; Bengtson et al., 2005; Kelly et al., 2010; Von Duyke et al., 2020; Vacquié-Garcia et al., 2021). Therefore, under the hypothesis that polar bears still-hunt more when seals spend more time in the water, bears should still-hunt during the day during winter, and during spring, bears should still-hunt during the night. However, opposite to this hypothesis Ware et al. (2020) found a peak in polar bear activity from 12:00–14:00 h during early winter, when seals haul-out at night, and a peak in bear activity at 24:00 h during the pupping season (i.e., April and May), when seals haul-out primarily during the day. The peak in activity we observed in January–March was 8 h later than Ware et al. (2020), however in line with Ware et al. (2020) we observed a night-time peak in activity in March–May followed by a highly variable timing of activity in June (Supplementary materials Fig. D1). The discrepancies in sleep and activity among studies may be due to population variability, interactions with season, or methodological differences (e.g., inability to differentiate between sleep and still-hunting).

As winter progressed, we observed a decrease in the low energy states (i.e., drift and ARS) and an increase in the higher energy state (i.e., olfactory search). The shift from low energy behaviors to high energy behaviors coincides with seals accessibility and may reflect corresponding changes in ideal hunting strategy from ambush predation to stalking predation (Messier et al., 1992; Ware et al., 2020; Vacquié-Garcia et al., 2021). Polar bears exhibit various hunting strategies with corresponding variation in movement and activity (Ware et al., 2020). During the first half of winter, seals spend the majority of their time in the water and rarely haul-out to the surface (Kelly et al., 2010; Von Duyke et al., 2020). Thus, access to ringed seals is primarily limited to breathing holes, which are mostly preyed on using still-hunting (Stirling, 1974). In April–May, ringed seals give birth and nurse their pups in subnivean lairs as the pups lack the thermal insulation to withstand the cold temperatures of Arctic waters and environment (Smith and Stirling, 1975). The subnivean lairs are visually inconspicuous, and polar bears rely on their sense of smell to locate them (Stirling and Latour, 1978; Togunov et al., 2017). Following the pupping season, from late May until the sea-ice melts, seals spend the majority of their time basking on the sea-ice surface to molt (Bengtson et al., 2005; Kelly et al., 2010; Von Duyke et al., 2020; Gryba et al., 2021). In addition, as the sea ice begins to thaw from mid-May, ringed seals no longer need to rely on breathing holes to surface, reducing the effectiveness of still-hunting (Stirling and McEwan, 1975; Messier et al., 1992). Access to seals is greatest during the pupping and molting season, which corresponds to the peak polar bear foraging period when bears enter hyperphagia (Stirling and Archibald, 1977; Messier et al., 1992; Pilfold et al., 2012; Reimer et al., 2019).

The increase in olfactory search during a period of increased prey availability aligns with the predictions of the marginal value theorem. The theorem states that a predator should exploit a patch until the energy intake rate drops to the average of the entire habitat, which would occur earlier during periods and regions with greater resource availability (Charnov, 1976). It is noteworthy that the marginal value theorem does not necessarily predict the specific mechanism that governs residency, but rather the departure time from a patch (Charnov, 1976). Residency time is an emergent property of discrete behaviors and finer-scale space use. For example, higher residency time can be produced by slower manifestation of some behavior (e.g., slower travel), longer time spent in slower behaviors (e.g., sleeping or nursing), or engaging in slow behaviors associated with patch use (e.g., ARS or feeding). We argue that a behavioral switch from still-hunting to active hunting is the behavioral mechanism that would lead to resource-linked residency time. A behavioral switch can also be viewed in terms of optimal foraging theory. For example, during periods of greater resource availability, animals can afford to engage in more energy-costly behaviors to maximize energy intake, and during periods of reduced resource availability, animals should switch to energy-conserving behaviors (Pyke et al., 1977). For example, during the summer fasting period when polar bears remain onshore, they exhibit the lowest annual activity in order to minimize energy expenditure (Parks et al., 2006; Whiteman et al., 2015; Togunov et al., 2017; Blanchet et al., 2020; Pagano et al., 2020). In contrast, bears with access to subsistence-harvested whales during the summer exhibit higher activity (Ware et al., 2017).

Most other polar bear populations exhibit a similar increase in activity coinciding with the seal pupping period (Stirling and Archibald, 1977; Smith, 1980; Messier et al., 1992; Amstrup et al., 2000; Blanchet et al., 2020; Ware et al., 2020). However, some contrasting patterns have been documented in the literature, which may be due to geographic or methodological differences. In areas of land-fast ice, polar bears increase frequency of still-hunting from spring to summer as warmer temperatures expose snow-covered breathing holes and promote still-hunting (Stirling and Latour, 1978). In areas of pack ice, warm temperatures promote the formation of open leads and impede still-hunting (Stirling and McEwan, 1975). Indeed, polar bear movement rates are higher over active ice than over consolidated ice (Ferguson et al., 2001; Blanchet et al., 2020). Parks et al. (2006) showed higher movement rates in winter compared to break-up in Hudson Bay, however they did not correct for sea ice motion, which is faster in winter (Yu et al., 2020) and can artificially inflate movement rates (Amstrup et al., 2000; Auger-Méthé et al., 2016).

The decrease in stationary behavior we observed in spring also coincides with the peak in polar bear breeding activity in March–May (Ferguson et al., 2001; Stirling et al., 2016; Ware et al., 2020). Polar bear breeding pairs are associated with reduced time walking and hunting compared to non-breeding pairs resulting in a corresponding range contraction (Stirling et al., 2016), which we expect to manifest as an increase in ARS-like movement. Assuming a typical weaning period between March and June of 2.25–2.5 year old cubs (Stirling and Latour, 1978; Ramsay and Stirling, 1988), four (one confirmed) of our 14 bears would have been alone and possibly breeding. We expect these bears to have inflated the frequency of ARS in spring, suggesting the observed increase in olfactory search may be conservative among non-breeding individuals. In addition, sea ice motion is faster earlier in the season (Yu et al., 2020), which may inflate movement rates, increase misclassification of drift as ARS or olfactory search, and further underestimate the seasonal decline in drift.

We hypothesized two effects of ice concentration on polar bear foraging. First, lower ice concentrations are more energetically costly to move through, wherein bears may have to travel longer distances to avoid swimming (Sahanatien and Derocher, 2012; Biddlecombe et al., 2021), which use significantly more energy than walking (Durner et al., 2011; Griffen, 2018). Second, ice concentration may influence the distribution and accessibility of seals. As described earlier, high ice concentrations may be more amenable to still-hunting as seal access to surface is more constrained and predictable, while at low ice concentrations, seals access to open water is greater and may promote active search and stalking hunt among polar bears (Stirling and McEwan, 1975; Messier et al., 1992). We observed an increase in the drift state as ice concentration increased in support of both the aforementioned hypotheses. In addition, the marginal value theorem predicts polar bears should spend less time in lower quality patches as the energy acquisition would reach the average of the entire habitat faster than in high quality patches (Charnov, 1976). Indeed, polar bears select for areas of high ice concentration, where they exhibit greater residency times, and lower movement rate in most studied populations (Ferguson et al., 2001; Durner et al., 2009; Pilfold et al., 2014; Laidre et al., 2015a; Lone et al., 2018; Bohart et al., 2021). Some studies identified an unexplained increase in activity with ice concentration (Ware et al., 2017; Blanchet et al., 2020), contrasting results that may be due to limited location data in low ice concentration in our research, different periods examined, or geographic variation.

There are open leads that encircle Hudson Bay and areas closer to shore tend to be more biologically productive (Stirling and Cleator, 1981; Henderson et al., 2021), resulting in ringed seals being more likely to remain in a resident behavior in shallower areas close to shore (Luque et al., 2014). Polar bear habitat selection with respect to distance from shore appears to depend on scale. At large scales (e.g., *>* 150 km), polar bears throughout the Arctic appear to select areas closer to shore (Durner et al., 2009; Laidre et al., 2015a; McCall et al., 2016; Johnson and Derocher, 2020). However, at a finer scale (e.g., *<* 150 km), polar bears appear to select for habitat further from shore, as the land-fast ice near shore may be less productive and preferentially used by subordinate individuals (Pilfold et al., 2014; Johnson and Derocher, 2020). In line with previous research, we observed a peak in the probability of being in the stationary drift state around 80 km, with olfactory search being more common closer to shore, and ARS increasing with distances *>* 80 km from shore. Thus, we suggest that there may be an optimal distance to shore rather than a simple monotonic increase or decrease as suggested by previous research. The selected distance to shore is likely specific to population, season, and demographic group. For example, in areas with a narrow continental shelf, such as the Beaufort and Greenland Seas, bears may remain closer to shore (Durner et al., 2009; Laidre et al., 2015a; Johnson and Derocher, 2020) compared to areas with a broad continental shelf (e.g., Hudson Bay and Laptev and Kara Seas; Durner et al., 2009; McCall et al., 2015, 2016). In areas with seasonal ice, bears may select areas close to shore to maintain proximity to summering grounds, which is supported by an increasing selection close to shore in break-up compared to winter (McCall et al., 2016). Competitive exclusion may also force subordinate individuals (e.g., females with cubs of the year) into lower quality habitat (Pilfold et al., 2014; McCall et al., 2015; Johnson and Derocher, 2020).

We predicted olfactory search would be most common at moderate wind speeds since still air and fast winds can impede olfaction (Aluja et al., 1993; Gibson and Torr, 1999; Togunov et al., 2017), and fast or cold winds may encourage polar bears to shelter in place (Harington, 1968) and deter seals from hauling out (Frost et al., 2004; Carlens et al., 2006; Gryba et al., 2021). However, as wind speed increased, we observed a decrease in drift and increase in ARS. This unexpected relationship between drift and ARS with respect to wind was likely an artifact of our ice motion correction model, which was trained on lower resolution dropped collars. Due to temporal averaging, the model may underestimate sea ice motion caused by wind, causing stationary behavior to be misclassified as ARS at high winds. There is a need for movement models that simultaneously predict and correct for sea ice motion and carry error in drift estimation forward into state classification.

### 5.2 State distribution

We observed variation in distribution associated with season, mean ice concentration, and bear age. Data from early winter were further from the summer refugia and had a larger range than data from late winter. Studies have found selection for areas closer to shore was stronger in late winter compared to early winter (Parks et al., 2006; McCall et al., 2016; Laidre et al., 2018). In contrast to our findings, Durner et al. (2019) identified a range expansion from winter into break-up in the Beaufort Sea that appeared to reflect a regime shift from stable multi-year sea ice to seasonal ice resulting in some of bears migrating south in the summer and some to range north following the retreating sea ice (Pilfold et al., 2016; Pagano et al., 2021).

The bear utilization distribution was greater in years with low ice concentration, which may be a response to habitat fragmentation that encourages bears to range further in search of quality habitat. Only 28% of the data points occurred in years with low ice concentration, suggesting that the observed area of the UD may be a conservative estimate. A similar negative relationship with sea ice concentration and home range size was found in several polar bear populations (Ferguson and Taylor, 1999; Amstrup et al., 2000; Hamilton et al., 2015; Durner et al., 2019; Pagano et al., 2021). However, other work showed home range to increase with an extended ice season and greater extent (Parks et al., 2006; Laidre et al., 2018). One possible explanation for the differences among studies may be that some populations experience habitat fragmentation (including Western Hudson Bay) while others experience habitat loss (Fahrig, 2003; Bonnell et al., 2018); habitat fragmentation may increase ranging in search of quality patches (e.g., Gardiner et al., 2019; Pagano et al., 2021), while habitat loss may lead to declines in extent as parts of the historic range become unavailable (e.g., Hinam and Clair, 2008; Stern and Laidre, 2016). If this hypothesis is correct, we predict that early stages of sea-ice decline (seasonal or interannual) may promote home range expansion, while latter stages of decline may cause home range decline.

Lastly, we observed that the utilization distribution of younger bears extended further from the summer refugia than older bears. One possible explanation is intraspecific competition, wherein older, dominant individuals may force subordinates into less optimal habitat through competitive exclusion or kleptoparasitism, a pattern observed between dominant males and solitary females relative to subordinate subadults and females with cubs in other polar bear populations (Stirling, 1974; Pilfold et al., 2014; McCall et al., 2015; Johnson and Derocher, 2020; Stirling et al., 2020; Florko et al., 2020b). An alternative explanation is age-specific navigational effectiveness, wherein younger bears have poorer navigation abilities on the dynamic drifting sea ice and move further from the summer refugia. Similarly, migrating passerines exhibit a “coastal effect” where younger, inexperienced passerines stray further from the optimal migration flyway compared to adults (Ralph, 1981). The lower navigation abilities hypothesis is supported in our observed behavior probability, which revealed that older individuals spent more time in low energy states. The marginal value theorem predicts that residency time should increase with patch quality, suggesting that older individuals may be more effective at locating higher quality patches (Charnov, 1976).

We incorporated orientation bias relative to wind to help differentiate behaviors with similar movement characteristics. This approach is limited when the external factor is spatiotemporally autocorrelated, as only the first few steps in a taxic behavior may display orientation bias, and once the desired angle is obtained, movement appears autocorrelated (Benhamou, 2006; Codling et al., 2008; Togunov et al., 2021). Behaviors of similar or shorter duration than the telemetry data or behavior transitions that occur between location data may lead to misclassification. In addition to state interpretation, it is not self-evident which behaviors are the most ecologically important. For example, olfactory search may represent optimal search strategy when conditions are favorable or a longer search for higher quality habitat (Iorio-Merlo et al., 2022). However, the diurnal presence of all three states throughout winter suggest that all three play an important ecological role. Investigating behaviors across seasons with resource variability may provide additional contrast to identify behavioral signatures associated with quality habitat (Cherry et al., 2016; Whiteman et al., 2015; Togunov et al., 2017; Ware et al., 2017; Pagano et al., 2020). In addition, future research should consider interactions between covariates; for example due to polar bear and seal seasonally varying diurnal movement and activity patterns (Parks et al., 2006; Von Duyke et al., 2020; Ware et al., 2020). Last, as analytical techniques continue to advance, programming at least some tags to a higher resolution (e.g., 1 h or 30 min) would enable research of finer-scale behaviors. Identifying baselines for behavioral time budgets may be increasingly important as environmental conditions continue to change.

### 5.3 Conclusion

Different behaviors have unique fitness and ecological consequences. Quantifying behavioral time-budgets and factors that promote different behaviors is key to understanding a species’ ecology. Remote tracking has elucidated much about polar bear habitat use (Laidre et al., 2022), however behavioral research has typically been limited to two states or less (e.g., Auger-Méthé et al., 2016; Ware et al., 2020, but see Pagano et al. 2017, 2020). Using advanced models that integrate wind data (Togunov et al., 2021), our study described previously undocumented circadian patterns in Western Hudson Bay polar bears, as well as behavioral variation with respect to season and ice concentration that appear to reflect a shift in foraging strategy in response to a change in prey availability (i.e., increase in haul-out behavior from early winter to the spring pupping and molting seasons). Last, we identified spatial patterns of distribution with respect to season, ice concentration, and bear age that may be indicative of habitat quality and competitive exclusion. Our findings expand on phenologic variation among polar bear populations that may be associated with regional or temporal variation in resource abundance or distribution. Due to the high degree of variation in ice dynamics throughout the Arctic, it is difficult to draw conclusions across populations (Amstrup et al., 2000; Klappstein et al., 2020). Given the circumpolar distribution of polar bears, each population is experiencing a different level of climate change related effects, stressing the need for population-specific research. The focal subpopulation of this paper – the Western Hudson Bay – is near the southern limits of the species’ range and is amongst the most affected by climate change. Our findings stress the importance of accounting for both behavior and the temporal and environmental factors that affect behavior. Failing to account for temporal factors that affect space use may obscure important habitat associations. For example, as polar bear behavior changes seasonally, resource selection functions that do not account for season may miss important patterns of habitat selection. Therefore, our methodology can help identify periods, locations, and environmental conditions that are associated with habitat quality to can help to better understand polar bear behavioral ecology and aid conservation.

## 6 Acknowledgments

Financial and logistical support of this study was provided by the B.C. Knowledge Development Fund Program, the Canada Foundation for Innovation (John R. Evans Leaders program) Canadian Association of Zoos and Aquariums, the Canadian Research Chairs Program, the Churchill Northern Studies Centre, Canadian Wildlife Federation, Care for the Wild International, Earth Rangers Foundation, Environment and Climate Change Canada, Hauser Bears, the Isdell Family Foundation, Kansas City Zoo, Manitoba Sustainable Development, Natural Sciences and Engineering Research Council of Canada, Parks Canada Agency, Pittsburgh Zoo Conservation Fund, Polar Bears International, Quark Expeditions, Schad Foundation, Sigmund Soudack and Associates Inc., Wildlife Media Inc., and World Wildlife Fund Canada.

