## Supplementary Materials for "Drivers of polar bear behavior, and the possible effects of prey availability on foraging strategy"

### A Drift correction

#### A.1 Drift correction methods

Togunov et al (2020) identified 20 drifting collars with locations within Hudson Bay between 2005–2015. To remove data-sparse periods, we segmented the tracks at any gaps larger than 24 hours and discarded segments with fewer than 16 locations. To obtain a regular time series for each track segment, we imputed missing locations by fitting a continuous-time correlated random walk model using the `crawl` package in `R` (Johnson and London, 2018; Johnson et al, 2008; RStudio Team, 2021). After filtering sparse locations and imputing missing locations (n = 725; 6.79%) each collar had a mean of $562 \pm 347$ GPS fixes (total = 10675).

Sea ice drift is driven by wind forcing, ocean currents, the Coriolis force, and internal ice stress. Due to the compressive strength of sea ice, local drift is typically affected by average force applied to it at a large scale. To allow for this, we summarized wind velocities at four buffers (0 km, 50 km, 100 km, 150 km). The buffer represents the radius around each GPS location from which wind velocities were extracted. The wind velocities of all raster cells that fell within the buffer were averaged to obtain the an estimate for the GPS point. For the buffer of 0 km, wind speed was estimated at the GPS location using bilinear interpolation. The buffer with the lowest AIC was 100 km and was used to predict drift. Wind velocities were obtained from the ERA5 meteorological reanalysis project, which provides hourly global analysis fields at a 31 km resolution (Hersbach et al, 2020) and were extracted using the `raster` package (Hijmans and van Etten, 2016).

The motion of the GPS tracks was described using two data streams: step length $l_t \in (0, \infty)$ (the distance between consecutive locations) and turning angle $\phi_t \in (-\pi, \pi]$ (change in bearing between consecutive steps), where $t \in 1, ..., T$ represents the time of the first GPS location in a step (Langrock et al, 2012; McClintock and Michelot, 2018). Step length $l_t$ and turning angle $\phi_t$ were modelled as slope-intercept functions of wind velocities. We fit Weibull and gamma distributions to step length and von Mises and wrapped Cauchy distributions to turning angle and selected the combination of distributions that had the lowest AIC (McClintock et al, 2020). Our final model for drift assumed that step length follows a Weibull distribution:

$$l_t \sim \text{Weibull}(\kappa_t^{(l)}, \lambda_t^{(l)}), \tag{1}$$

where $\kappa_t^{(l)} > 0$ is the shape parameter and $\lambda_t^{(l)} > 0$ is the scale parameter of the step length (McClintock et al, 2020). The shape and scale parameters were each defined as slope-intercept functions of wind speed

$r_t \geq 0$ as follows:

$$\ln(\kappa_t^{(l)}) = \beta_1 + \beta_2 r_t \tag{2}$$

$$\ln(\lambda_t^{(l)}) = \beta_3 + \beta_4 r_t, \tag{3}$$

where is $\beta_{\{1,2,3,4\}} \in \mathbb{R}$ are the intercept and slope coefficients to be estimated.

According to AIC, our best model defined turning angle $\phi_t$ using the von Mises distribution. Due to the Coriolis effect, sea ice drift is approximately -20°relative to the wind direction in the northern hemisphere (Tschudi et al, 2010). However, the precise angle depends on a number of factors and was estimated closer to -15°in Hudson Bay (Togunov et al, 2021). To allow for drift toward an unknown angle of bias relative to wind, we modelled the mean turning angle as a trade-off between bias parallel to the direction of the wind and bias perpendicular to wind. Specifically, mean turning angle $\mu_t^{(\phi)}$ was assumed to follow a menotactic BRW circular-circular von Mises regression model (Rivest and Duchesne, 2016; Togunov et al, 2021):

$$\begin{aligned} \mu_t^{(\phi)} = \operatorname{atan2}(&\alpha_1 \sin\psi_t + \alpha_2 \cos\psi_t, \\ &\alpha_1 \cos\psi_t - \alpha_2 \sin\psi_t), \end{aligned} \tag{4}$$

where $\psi_t$ is the direction of wind at $t$ relative to the movement bearing at $t-1$. We hypothesized that the concentration of turning angle around $\mu_t^{(\phi)}$ would increase as wind speeds increased. At higher wind speeds, the force is greater and the direction can be predicted more accurately. To capture this, we modeled turning angle concentration as a slope-intercept function of wind speed following:

$$\ln(\kappa_t^{(\phi)}) = \beta_5 + \beta_6 r_t. \tag{5}$$

After fitting the BCRW drift model to the dropped collar data, we predicted the drift speed and direction given the estimated ECMWF wind vectors. Shape and scale parameters of step length were estimated using Eq. 2 and Eq. 3, which were used to estimate the latent drift speed $\hat{l}_t$ as the mode of Weibull step length following:

$$\hat{l}_t = \begin{cases} 0 & \text{if } \hat{\kappa}_t^{(l)} \leq 1 \\ \hat{\lambda}_t \left( \dfrac{\hat{\kappa}_t^{(l)} - 1}{\hat{\kappa}_t^{(l)}} \right)^{1/\hat{\kappa}_t^{(l)}} & \text{if } \hat{\kappa}_t^{(l)} > 1. \end{cases} \tag{6}$$

We used the estimated mode drift speed for the correction as it minimized the median drift-corrected step length $l_t^{(c)}$ compared to the mean or median of the Weibull distribution.

Mean direction $\hat{\mu}_t^{(\phi)}$ of drift was predicted using Eq. 4. Next, we estimated the drift-corrected paths by subtracting the estimated drift vectors from the observed drift vectors following:

$$\begin{aligned} \hat{u}_t &= \hat{l}_t \cos \hat{\mu}_t^{(\phi)} \\ \hat{v}_t &= \hat{l}_t \sin \hat{\mu}_t^{(\phi)}, \end{aligned} \tag{7}$$

where $\hat{u}_t \in \mathbb{R}$ and $\hat{v}_t \in \mathbb{R}$ are the estimated east-to-west and south-to-north components of the drift-corrected track. The estimated drift-corrected step length $\hat{l}_t^{(c)}$ was calculated following:

$$\hat{l}_t^{(c)} = \sqrt{\hat{u}_t^2 + \hat{v}_t^2}. \tag{8}$$

Next, we estimated the drift-corrected trajectory of the track using the cumulative sum of the estimated $u_t$ and $v_t$ components following:

$$\begin{aligned} \hat{x}_t^{(c)} &= \begin{cases} [h]0 & \text{if } t = 1 \\ \sum_{i=1}^{t-1} \hat{u}_i & \text{Otherwise} \end{cases} \\ \hat{y}_t^{(c)} &= \begin{cases} 0 & \text{if } t = 1 \\ \sum_{i=1}^{t-1} \hat{v}_i & \text{Otherwise.} \end{cases} \end{aligned} \tag{9}$$

where $\hat{x}_t^{(c)}$ and $\hat{y}_t^{(c)}$ are the estimated x and y coordinates of the drift-corrected track, respectively.

To test that we did not over-fit the data, we evaluated the drift correction BCRW performance in reducing the apparent speed of dropped collars by fitting it to a random subset of 10% of the dropped collar tracks and correcting for it in the entire dropped collar data set. When correcting for drift in the bear tracks, we fit the drift correction BCRW to the entire dropped collar data-set, and used the estimated coefficients and ECMWF wind to predict and correct for drift along the bear tracks.

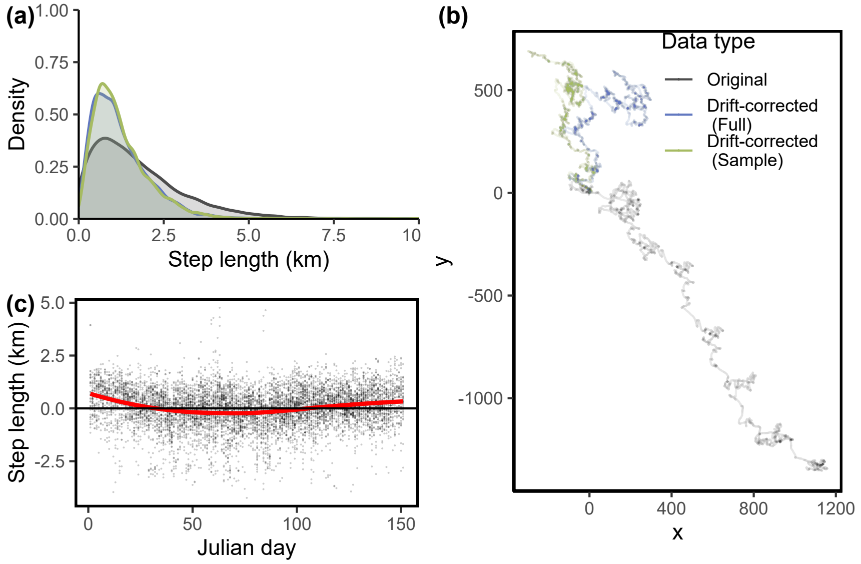


Figure A1: Drift correction: (a) Step length of dropped collars before and after drift correction; (b) cumulative path of dropped collars before and after drift correction; (c) residual plot of step length for drift corrected dropped collars sorted by ordinal date. Grey represents original data, blue represents drift-corrected data based on BCRW fit to the full data set, and green represents drift-corrected data based on BCRW fit to a 10% random sample of dropped collar tracks.

#### A.2 Drift correction results

We expect that applying the drift correction would mostly reduce the movement of the drop collar close to zero. When fit to the full dropped collar data set, the drift correction reduced the mean step length by 29% from 1.76 km to 1.25 km (Fig A1). When fit to a 10% random sample of dropped collar tracks, the drift correction reduced the mean step length by 28% to 1.26 km. (Fig. A1). The drift corrected tracks still appear to have a remaining directional bias, however it is less than the original track (Fig. A1). There appears to be some residuals with respect to day of the year (Fig. A1C), however, including day of the year in the drift correction BCRW yielded only marginal model improvement with respect to reduction of step length (data not shown).

### B Ice motion-based drift correction

#### B.1 Drift correction methods

We tested using sea ice motion vectors produced by the National Snow and Ice Data Center (NSIDC; Tschudi et al, 2019, 2020) to correct for ice motion in the dropped collars as in Appendix A.

All the same procedures as described in Appendix A.1 were followed with the exception of the definition of step length $l_t \in (0, \infty)$ and turning angle $\phi_t \in (-\pi, \pi]$ being defined in terms of sea ice drift rather than wind velocities. Specifically, the shape and scale parameters of step length were defined as slope-intercept functions of NSIDC ice motion speed $r_t^{(NSIDC)} \geq 0$ as follows:

$$\ln(\kappa_t^{(l)}) = \beta_1 + \beta_2 r_t^{(NSIDC)} \tag{10}$$

$$\ln(\lambda_t^{(l)}) = \beta_3 + \beta_4 r_t^{(NSIDC)}, \tag{11}$$

where is $\beta_{\{1,2,3,4\}} \in \mathbb{R}$ are the intercept and slope coefficients to be estimated.

Mean turning angle $\mu_t^{(\phi)}$ was assumed to be a BRW circular-circular von Mises regression model (Rivest and Duchesne, 2016):

$$\begin{aligned} \mu_t^{(\phi)} = \text{atan2}(&\alpha_1 \sin \psi_t^{(NSIDC)}, \\ &\alpha_1 \cos \psi_t^{(NSIDC)}), \end{aligned} \tag{12}$$

where $\psi_t^{(NSIDC)}$ is the direction of NSIDC ice motion at $t$ relative to the movement bearing at $t-1$. In addition, we modeled turning angle concentration as a slope-intercept function of NSIDC ice motion speed following:

$$\ln(\kappa_t^{(\phi)}) = \beta_5 + \beta_6 r_t^{(NSIDC)}. \tag{13}$$

After fitting the BCRW drift model to the dropped collar data, we predicted the drift speed and direction given the estimated NSIDC ice motion vectors. Shape and scale parameters of step length were estimated using Eq. 10 and Eq. 11 and were used to estimate the latent drift speed $\hat{l}_t$ using eq. 6. Mean drift direction $\hat{\mu}_t^{(\phi)}$ of drift was predicted using Eq. 12. Finally, we estimated the drift-corrected paths, step lengths, and trajectory as described in Appendix A.1 using eq. 7, eq. 8, and eq. 9, respectively

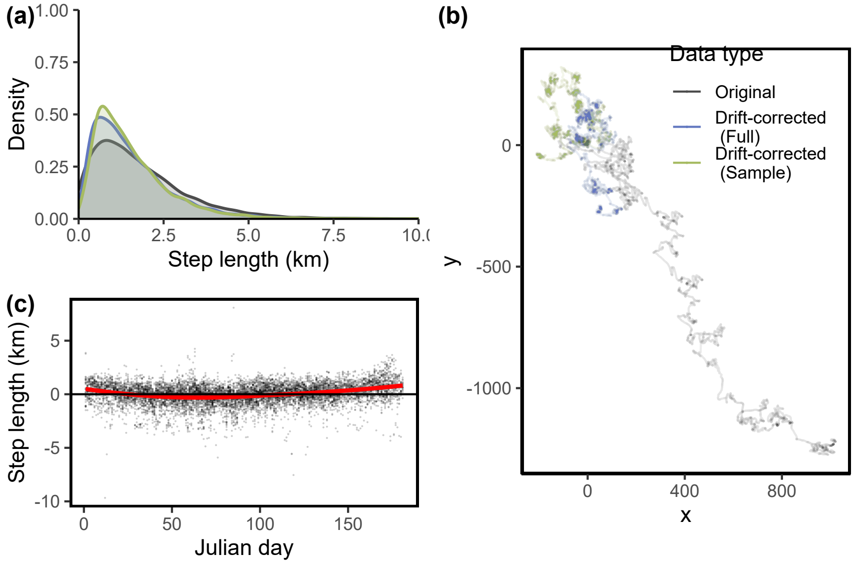


Figure B1: Drift correction: (a) Step length of dropped collars before and after drift correction using NSIDC sea ice motion vectors; (b) cumulative path of dropped collars before and after drift correction; (c) residual plot of step length for drift corrected dropped collars sorted by ordinal date. Grey represents original data, blue represents drift-corrected data based on BCRW fit to the full data set, and green represents drift-corrected data based on BCRW fit to a 10% random sample of dropped collar tracks.

#### B.2 Drift correction results

When fit to the full dropped collar data set, the drift correction reduced the mean step length by 13% from 1.81 km to 1.57 km; Fig. B1). When fit to a 10% random sample of dropped collar tracks, the drift correction reduced the mean step length by 13% from 1.81 km to 1.57 km (Fig. B1).

### C Pseudo-design matrices

Pseudo-design matrix for step length in `momentuHMM`:

$$\begin{array}{c} \\ \mu^{(l)}_{D,t} \\ \mu^{(l)}_{ARS,t} \\ \mu^{(l)}_{O^{(L)},t} \\ \mu^{(l)}_{O^{(R)},t} \\ \sigma^{(l)}_{D,t} \\ \sigma^{(l)}_{ARS,t} \\ \sigma^{(l)}_{O^{(L)},t} \\ \sigma^{(l)}_{O^{(R)},t} \end{array} \begin{array}{c} \begin{array}{ccccccc} \beta_{1,D} & \beta_{2,D} & \beta_{1,ARS} & \beta_{1,O^{(L,R)}} & \sigma_{1,D} & \sigma_{1,ARS} & \sigma_{1,O^{(L,R)}} \end{array} \\ \begin{pmatrix} 1 & r^{(tide)}_t & 0 & 0 & 0 & 0 & 0 \\ 1 & r^{(tide)}_t & 1 & 0 & 0 & 0 & 0 \\ 1 & r^{(tide)}_t & 1 & 1 & 0 & 0 & 0 \\ 1 & r^{(tide)}_t & 1 & 1 & 0 & 0 & 0 \\ 0 & 0 & 0 & 0 & 1 & 0 & 0 \\ 0 & 0 & 0 & 0 & 0 & 1 & 0 \\ 0 & 0 & 0 & 0 & 0 & 0 & 1 \\ 0 & 0 & 0 & 0 & 0 & 0 & 1 \end{pmatrix} \end{array} \tag{14}$$

The column correspond to the coefficients estimated using maximum likelihood by the model, and the rows correspond to the movement parameters: step length mean ($\mu^{(l)}_{S,t}$) and standard deviation ($\sigma^{(l)}_{S,t}$). $r^{(tide)}_t$ represents tidal current speed in m s$^{-1}$. We ensured that $\mu^{(l)}_{D,t} < \mu^{(l)}_{ARS,t} < \mu^{(l)}_{O^{(L,R)},t}$ using the `workBounds` argument in `fitHMM` by constraining $\beta_{1,ARS}$ and $\beta_{1,O^{(L,R)}}$ to be $> 0$, see McClintock and Michelot (2018) for details.

Pseudo-design matrix used for turning angle:

$$\begin{array}{c} \\ \mu^{(\phi)}_{D,t} \\ \mu^{(\phi)}_{ARS} \\ \mu^{(\phi)}_{O^{(L)},t} \\ \mu^{(\phi)}_{O^{(R)},t} \\ \kappa^{(\phi)}_{D} \\ \kappa^{(\phi)}_{ARS} \\ \kappa^{(\phi)}_{O^{(L)}} \\ \kappa^{(\phi)}_{O^{(R)}} \end{array} \begin{array}{c} \begin{array}{cccccc} \alpha_{1,D} & \alpha_{1,O^{(L,R)}} & \alpha_{2,O^{(L,R)}} & \kappa_{1,D} & \kappa_{1,ARS} & \kappa_{1,O^{(L,R)}} \end{array} \\ \begin{pmatrix} \texttt{angleFormula}(\psi^{(tide)}_t, r^{(tide)}_t) & 0 & 0 & 0 & 0 & 0 \\ 0 & 0 & 0 & 0 & 0 & 0 \\ 0 & \psi^{(wind)}_t & \psi^{(wind)}_t + 90^\circ & 0 & 0 & 0 \\ 0 & \psi^{(wind)}_t & \psi^{(wind)}_t - 90^\circ & 0 & 0 & 0 \\ 0 & 0 & 0 & 1 & 0 & 0 \\ 0 & 0 & 0 & 0 & 1 & 0 \\ 0 & 0 & 0 & 0 & 0 & 1 \\ 0 & 0 & 0 & 0 & 0 & 1 \end{pmatrix} \end{array} \tag{15}$$

`angleFormula` is a special function that can be included in design matrices to model circular parameters (e.g., mean turning angle $\mu^{(\phi)}_{D,t}$) as a circular-circular regression function of the a directional covariate and its relative strength - in this instance, tidal current direction, $\psi^{(tide)}_t$, and speed, $r^{(tide)}_t$ (Rivest and Duchesne, 2016; McClintock and Michelot, 2018). $\psi^{(wind)}_t$ represents the direction of wind at time $t$ relative to the animal bearing at time $t-1$.

### D Diurnal and seasonal state Frequency

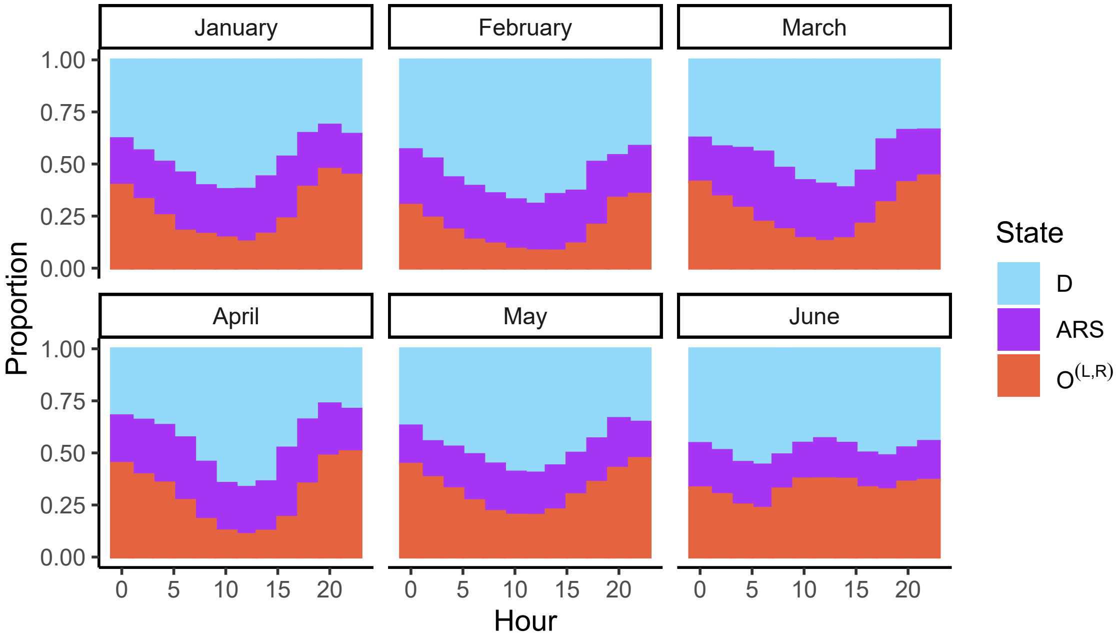


Figure D1: Proportion of states (drift $D$, area-restricted search $ARS$, and olfactory search $O^{(L,R)}$) depending on month and hour. States are based on Viterbi-decoded states.

### E State distribution map

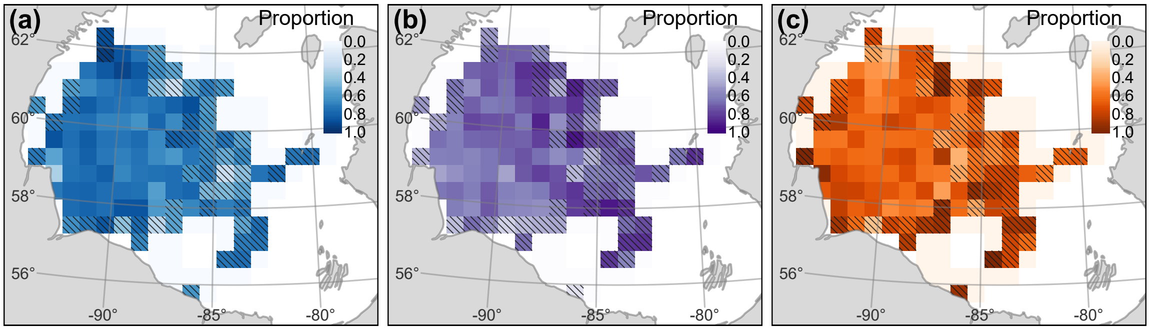


Figure E1: Distribution of predicted states. (a) proportion of bear-days for the drift state $D$, (b) proportion of bear-days for the area-restricted search state $ARS$, and (c) proportion of bear-days for the olfactory search state $O$. Cells with $< 7$ bear-days were not plotted and cells with $< 21$ bear days were hashed.
